# Cancer-associated fibroblasts reveal aberrant DNA methylation across different types of cancer

**DOI:** 10.1101/2024.02.02.578626

**Authors:** Marco Schmidt, Tiago Maié, Ivan G. Costa, Wolfgang Wagner

## Abstract

Cancer-associated fibroblasts (CAFs) are critical components of the tumor microenvironment. Several studies demonstrated molecular differences between CAFs and normal tissue-associated fibroblasts (NAFs). In this study, we isolated CAFs and NAFs from liver tumors and analyzed their DNA methylation profiles. A subset of the CAFs exhibited aberrant DNA methylation, which was also reflected on gene expression level. The DNA methylation at liver-CAF-specific CG dinucleotides (CpGs) was associated with survival in liver cancer data. An integrative analysis with public datasets of CAF *versus* NAF in different cancer types, including lung, prostate, esophagus, and gastric cancer, revealed overlapping epigenetic aberrations. CpGs with common aberrations in DNA methylation included cg09809672 (*EDARADD*), cg07134930 (*HDAC4*), and cg05935904 (intergenic). Aberrant DNA methylation at these sites was associated with prognosis in several cancer types. Thus, activation of CAFs by the tumor environment is associated with characteristic epigenetic modifications that could be used as biomarkers for disease stratification.

## Introduction

Cancer-associated fibroblasts (CAFs) modulate the microenvironment and are important for tumor development, metastasis, and resistance to therapy (Ping et al., 2021). In contrast to cancer cells, CAFs usually do not harbor genetic mutations – rather, their activation is reflected in various epigenetic changes (Kehrberg et al., 2023). A bottleneck in the molecular characterization of CAFs is that fibroblasts generally resemble complex cell populations with striking inter- and intra-organic heterogeneity (Muhl et al., 2020). The heterogeneity of CAFs is further complicated by diverse cellular origins that can be recruited and activated during tumorigenesis, including normal-tissue associated fibroblasts (NAFs), bone marrow-derived mesenchymal stromal cells (MSCs), stellate cells, epithelial cells, and endothelial cells (Zhao et al., 2023). This heterogeneity makes it difficult to precisely define CAFs at the molecular level and may lead to a different mechanistic contribution of CAF subtypes to cancer pathophysiology (Bu et al., 2019).

The identification of suitable molecular biomarkers for CAFs remains a major challenge (Ping et al., 2021). Several “CAF markers” - including alpha smooth muscle actin (αSMA), vimentin, and fibroblast activation protein alpha (FAP) - have been proposed, but their expression varies between different cancer types and CAF subpopulations (Nurmik et al., 2020). To date, no single biomarker has been identified that can reliably discern CAFs and normal fibroblasts in a given tumor (Chen et al., 2021). Integration of transcriptomic data for multiple markers into “CAF scores” has provided prognostic information for different cancer types (Zou et al., 2022), and single cell RNA-sequencing data have revealed gene expression signatures of independent CAF subtypes that are even common to different cancer types (Galbo et al., 2021). It therefore seems conceivable that there are also epigenetic signatures of CAFs that are common to different tumor types.

Changes in DNA methylation (DNAm) are tightly controlled in a very consistent manner during normal cellular differentiation. Our group and others have already shown that DNAm at single CG dinucleotides (CpGs) can provide reliable biomarkers for specific cell types, e.g. fibroblasts and MSCs (de Almeida et al., 2016; Hubens et al., 2023; Schmidt et al., 2020). CAFs have also been shown to have different DNAm patterns for different cancer types (Kehrberg et al., 2023; Lawrence et al., 2020), and it has been proposed that changes in the methylome of CAFs represent a novel epigenetic feature of the cancer microenvironment that could provide therapeutically relevant biomarkers (Pidsley et al., 2018). However, so far it has been largely unclear if whether such DNAm alterations are also consistent across CAFs in different cancer types (Kehrberg et al., 2023).

In this study, we demonstrate that CAFs from liver tumors exhibit aberrant DNAm compared to fibroblasts from adjacent tissue. A comprehensive comparison with CAF-associated DNAm in lung cancer, esophageal carcinoma, prostate carcinoma and gastric cancer demonstrated an overlap, which could even be of prognostic importance.

## Results

### Aberrant DNA methylation in cancer-associated fibroblasts from liver tumors

Fibroblasts were isolated from cancer (CAFs) and from tumor-free adjacent tissue (NAFs) of hepatocellular carcinoma (HCC) or liver metastasis of colorectal/anal cancer (in total n = 11 CAF/NAF pairs). All cell preparations showed typical fibroblastoid morphology and surface marker expression (CD14^-^, CD29^+^, CD31^-^, CD34^-^, CD45^-^, CD73^+^, CD90^+^, and CD105^+^). In addition, immunostaining demonstrated that they were positive for vimentin, negative for pan-cytokeratin, and heterogeneous for alpha-smooth muscle actin (α-SMA), indicating that all cell preparations could be classified as fibroblasts (Supplemental Figure S1). The DNA methylation profiles were then analyzed with Illumina MethylationEPIC BeadChips. For orientation, we initially selected 2,134 CpG sites with at least 20% difference in mean DNA methylation in CAFs *versus* NAFs. Hierarchical clustering of these CpGs indicated that the CAF profiles can be categorized in two groups, which we subsequently refer to as CAF^high^ and CAF^low^ (Figure 1A). The CAF^high^ samples were also clearly separated in a multidimensional scaling (MDS) plot of the 10,000 most variable CpG sites (Figure 1B). This suggests that CAF-specific epigenetic aberrations vary between samples, and we therefore focused particularly on the epigenetic differences between CAF^high^ and NAFs.

**Figure 1:**
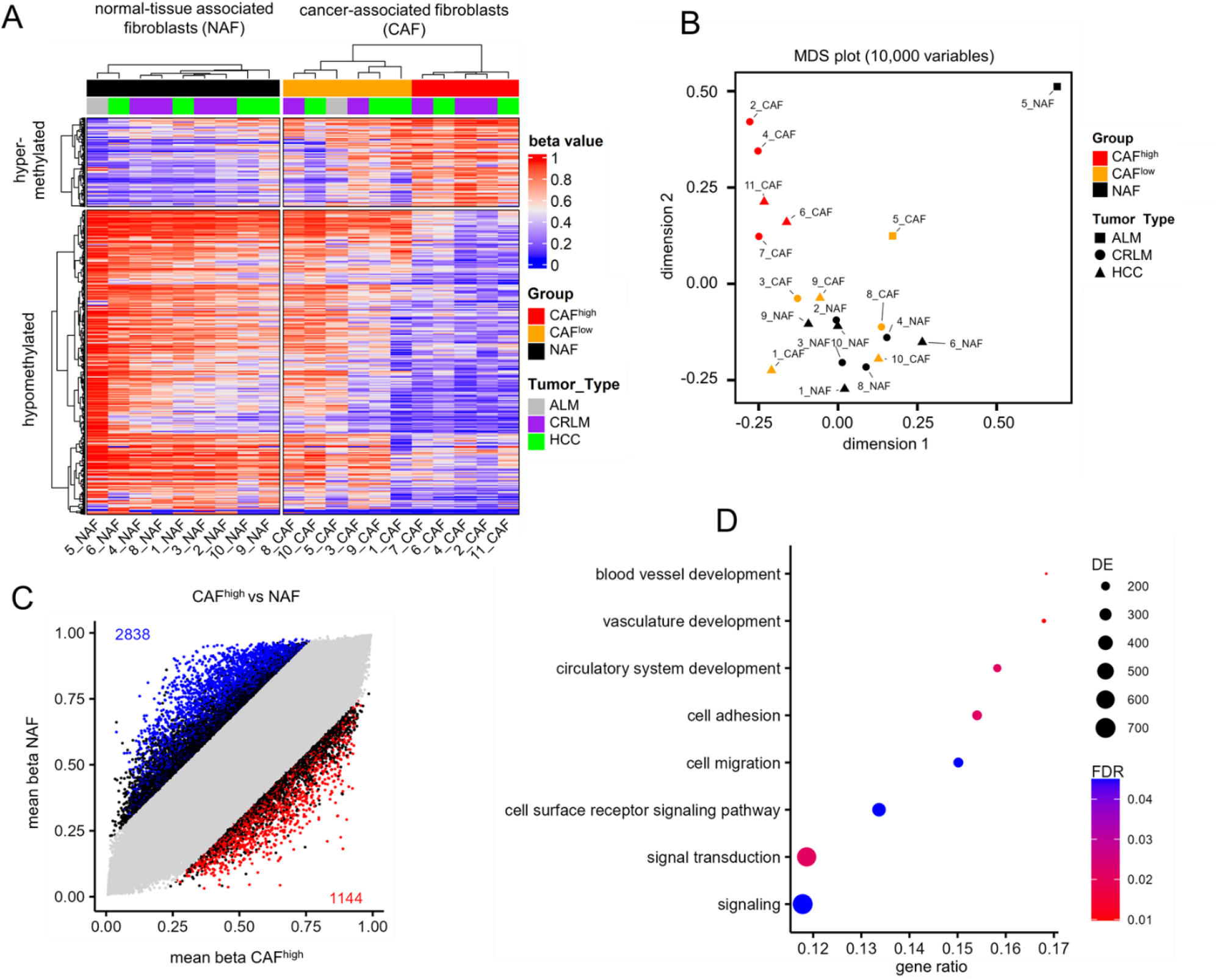
Aberrant DNA methylation in fibroblasts of liver cancer. (A) Liver fibroblasts were isolated from cancer tissue (CAFs) or from tumor-free neighboring tissue (NAFs) and analyzed on EPIC bead chips. The heatmap depicts DNA methylation at 2,134 CpGs with at least 20% mean methylation difference between NAFs and CAFs. Hierarchical clustering showed two groups of CAFs, which were referred to as CAF^low^ (orange) and CAF^high^ (red). (B) The multidimensional scaling (MDS) plot of the 10,000 most variable CpGs showed that CAF^high^clustered apart from CAF^low^ and NAFs. (c) Scatterplot comparing the mean beta values of the CAF^high^ group *versus* NAFs. Significant differentially methylated CpGs are highlighted (mean DNAm difference >20%; limma adjusted p-values < 0.05). (D) Gene ontology enrichment („biological process”) of CpG sites with significant differential DNAm between CAF^high^ and NAFs (DE = number of differentially methylated genes, FDR = false discovery rate).

Differential methylation analysis (> 20% difference in mean methylation; limma adjusted p-values < 0.05) revealed 2,838 significantly hypomethylated and 1,144 hypermethylated CpGs (Figure 1C; Supplemental Figure S2A,B; Supplemental Table S1). Gene ontology analysis revealed that aberrant DNA methylation was significantly enriched in categories associated with angiogenesis, cell migration, and signaling (Figure 1D). The differently methylated sites appeared to be located in gene bodies rather than promoter regions (Supplemental Figure S2C). The most significantly hypomethylated region was in the collagen type I alpha 1 chain (*COL1A1*) gene, a gene that has been used in the literature as a gene expression marker for a subset of CAFs (Bhattacharjee et al., 2021; Galbo et al., 2021; Zou et al., 2022), and the top hypermethylated region was in the gap junction protein alpha 4 (*GJA4*; Supplemental Figure S2D-E). These differentially methylated regions could therefore contribute to the activation of CAFs in liver tumors.

### Cancer-associated fibroblast reveal aberrant gene expression that reflects epigenetic aberrations

To better understand if the aberrant DNAm in CAFs is also reflected at the gene expression level, we performed RNA sequencing on seven of the NAF/CAF pairs. Analogous to the DNA methylation analysis, we initially selected at least fourfold difference in gene expression between NAFs and CAFs. This analysis resulted in the same separation of CAF^high^ and CAF^low^ samples as previously observed (Figure 2A-B). Further analysis of differential gene expression between the CAF^high^ and NAF group revealed 525 upregulated and 359 downregulated genes (Figure 2C; Supplemental Figure S3). Amongst the most significant upregulated genes were plakophilin-2 (*PHK2*), which has been described as a Wnt/β-catenin target in colon cancer CAFs (Niell et al., 2018), and TIMP metallopeptidase inhibitor 3 (*TIMP3*), which was also highly upregulated in ovarian cancer CAFs (Zeng et al., 2022).

**Figure 2:**
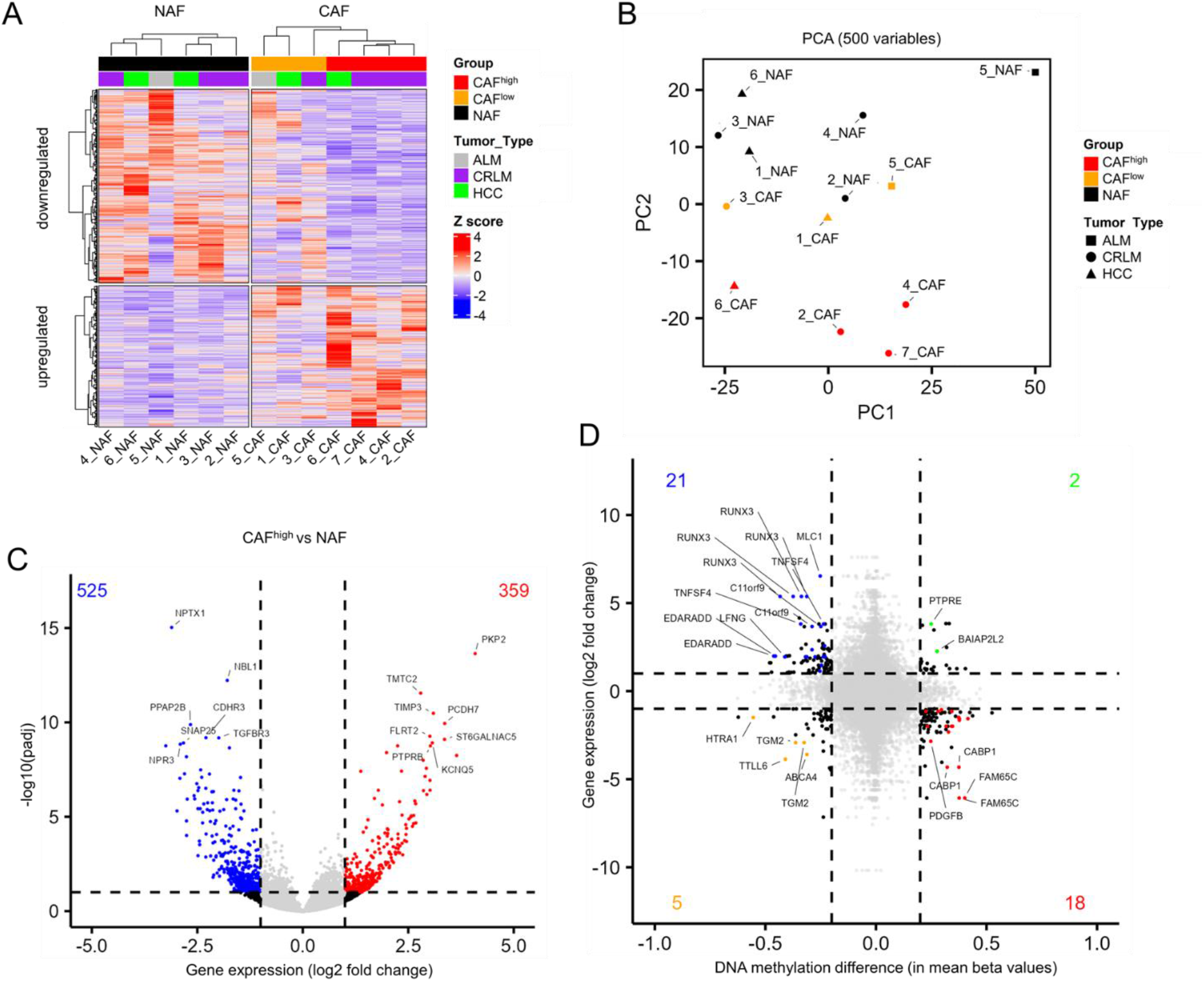
Gene expression differences between CAFs and NAFs from liver. (A) Heatmap of 891 genes with at least 4-fold expression difference between the groups of NAFs and CAFs. Hierarchical clustering showed the same classification of CAF^low^ (orange) and CAF^high^ (red), as observed for DNAm. (B) Principal component analysis of the 500 most variable genes. (c) Volcano plot comparing the CAF^high^ group with NAFs. Highlighted are significantly different expressed genes (adjusted p-values < 0.1). (D) Differential gene expression was compared with differential methylation in CAF^high^ *versus* NAFs. Each CpG site in promoter regions (TSS1500 and TSS200) was paired with the associated genes. Highlighted are significantly differentially expressed and methylated genes.

We then analyzed whether the aberrant DNA methylation in the CAF^high^ fraction was also reflected in corresponding changes in gene expression. In general, hypomethylation was associated with higher gene expression, and the opposite was also true (Figure 2D). Interestingly, EDAR Associated Via Death Domain (*EDARADD*) was among the genes that showed high expression and hypomethylation. This gene has previously been independently described as differentially methylated and expressed in CAFs from lung and prostate cancer (Lawrence et al., 2020; Vizoso et al., 2015). In addition, the Runt-related transcription factor 3 (*RUNX3*) was amongst the highest expressed genes, as previously described for tumor-supporting CAFs from breast cancer (Koyama et al., 2023).

### Selection of candidate CpGs for cancer-associated fibroblasts in liver tumors

A potential epigenetic biomarker for CAFs should have consistent DNAm patterns in other cell types. We therefore searched for CpG sites with distinct DNAm levels between CAFs and other cell types from liver, including liver cancer cells. To this end, we compiled a dataset of in-house and public methylation data with a wide range of different cell types (Supplemental table S2). CAFs, NAFs and fibroblasts were close to each other in the MDS plot, while hematopoietic cells and cancer cell lines/cholangiocyte cancer cells were separated into distinct clusters (Figure 3A). Notably, methylome of CAFs was closely related to that of fibroblasts, supporting the notion that CAFs are of fibroblastoid origin.

**Figure 3:**
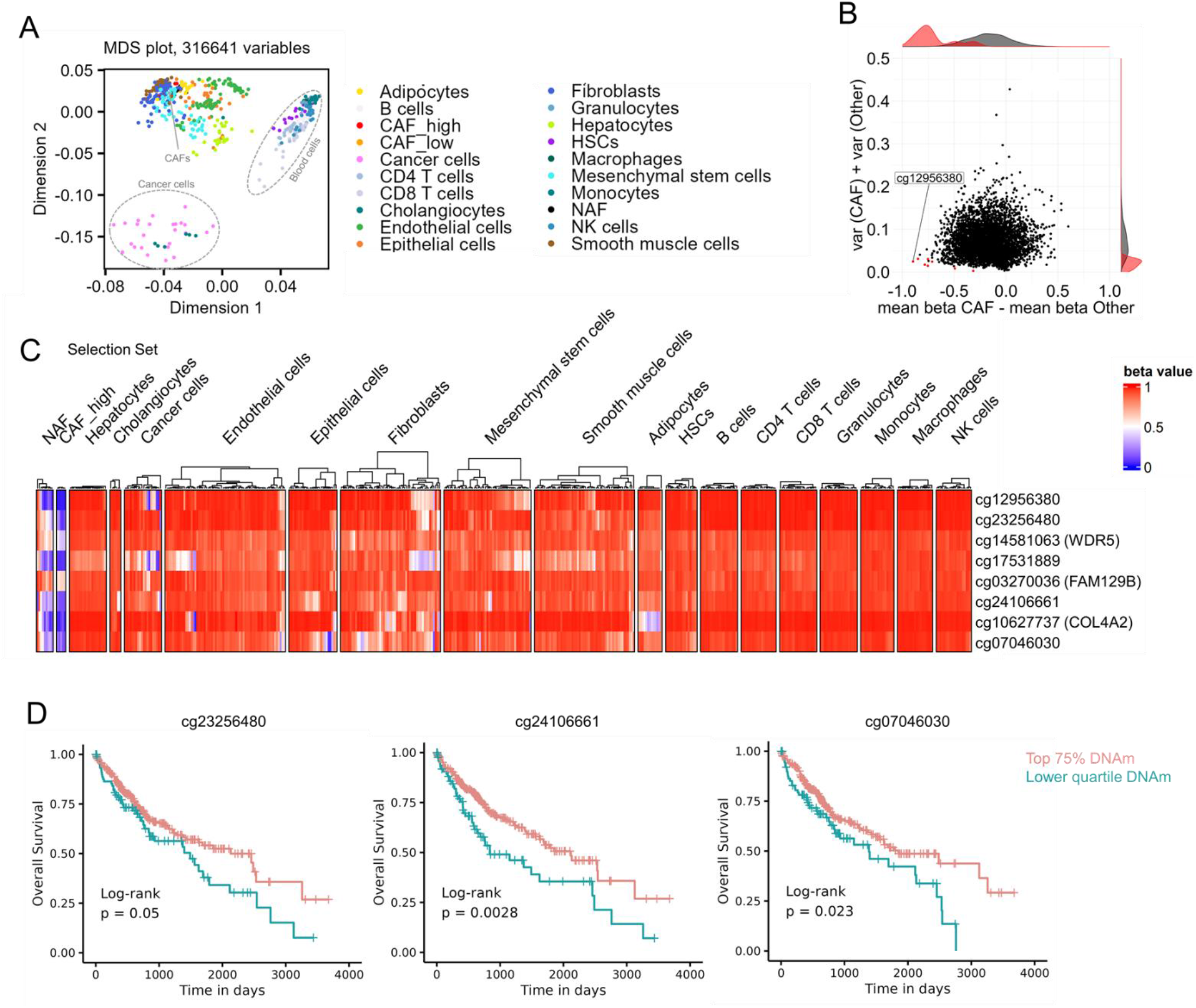
Selection of potential DNA methylation biomarkers for CAFs in liver cancer. (A) Multidimensional scaling (MDS) plot of DNA methylation profiles (316641 CpGs) in NAFs, CAFs with public datasets of various other cell types. Our NAFs and CAFs clustered closely to fibroblasts of other studies. (B) The selection of candidate CpGs was performed with CimpleG (Maie et al., 2023) on a reduced number of CpGs, that showed at least 20% mean methylation difference between NAFs and CAFs. (C) Heatmap of DNAm of eight candidate CpGs that were selected to discern CAFs from other cell types. The results of the selection dataset are depicted here. (D) To investigate if DNA methylation at these eight candidate CpGs is associated with overall survival, we used the TCGA data of hepatocellular carcinoma (Cancer Genome Atlas Research Network. Electronic address and Cancer Genome Atlas Research, 2017). Kaplan-Meier analysis of the 25^th^ percentile of patients with the lowest DNA methylation at these sites *versus* other patients revealed significant results for three CpGs (cg24106661; cg07046030; and cg23256480).

To identify CAF-associated CpGs, we divided the datasets into a selection set (including CAF^high^ samples) and test set (with CAF^low^ samples). We used CimpleG with a preselected number of CpGs (>20% mean difference between NAFs and CAFs) to select candidate CpGs (Figure 3B) (Maie et al., 2023). We exemplified the top eight candidate CpGs, all of which were hypomethylated in the CAF^high^ samples and consistently methylated in other cell types in the selection set (Figure 3C) as well as in the test set (Supplemental Figure S4A). We then tested whether the eight candidate CpGs for CAFs in liver tissue would also reveal different DNAm in hepatocellular cancer compared to normal liver tissue (Supplemental Figure S4B). Indeed, in tumor samples the candidate CpGs showed overall lower methylation compared to normal liver tissue.

To assess whether the DNAm pattern in these CAF-associated CpGs is also indicative for prognosis in liver cancer, we compared the 25^th^ percentile of patients with the lowest methylation at these sites with other patients. Three of the eight CpGs showed significant association with survival (cg24106661; cg07046030; and cg23256480; Figure 3D). Thus, lower DNA methylation at these CpGs may reflect a higher proportion of CAFs contributing to shorter long-term survival in hepatocellular carcinoma.

### Aberrant DNA methylation of cancer-associated fibroblasts in different types of cancer

To investigate if the epigenetic aberrations overlap in different cancer types, we used all at the time publicly available Illumina BeadChip profiles of CAFs and NAFs: from non-small cell lung cancer (Vizoso et al., 2015), prostate cancer (Lawrence et al., 2020), adenocarcinomas of the stomach and esophagus (Najgebauer et al., 2019), and three other datasets with CAFs from gastric cancer (Maeda et al., 2020; Najgebauer et al., 2019) (Supplemental Table S2). The samples were mainly grouped according to the tissue of origin in dimensions 1 and 2 in the multidimensional scaling (MDS; Figure 4A). Notably, dimension 4 of the MDS analysis separated CAF from NAF samples across all five different tissues, indicating that there may indeed be overlapping epigenetic differences (Figure 4B).

**Figure 4:**
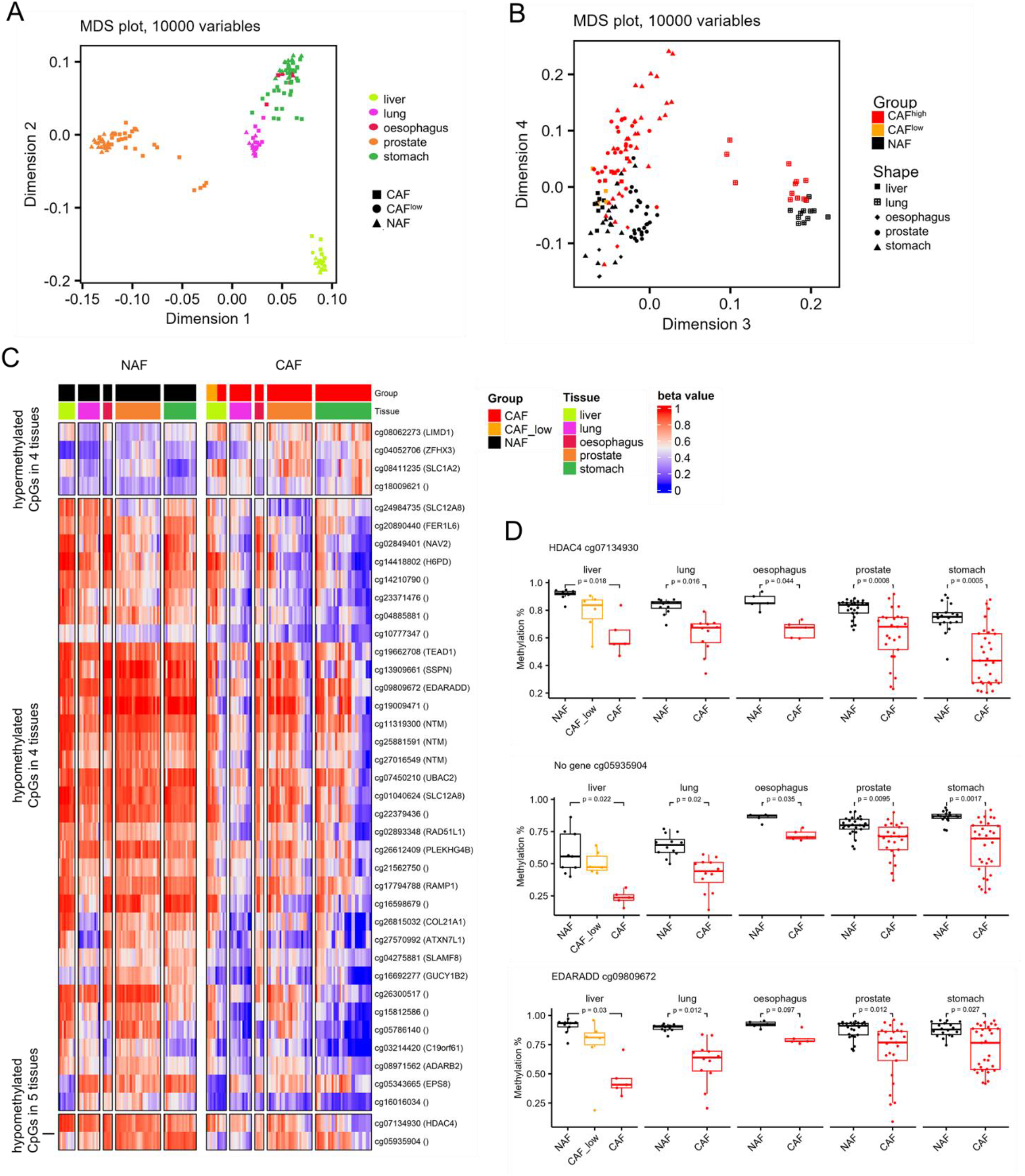
Aberrant DNA methylation in fibroblasts associated with various types of cancer. (A, B) Multidimensional scaling (MDS) plot of the 10,000 most variable CpGs in the dataset containing NAFs and CAFs from lung (GSE68851) (Vizoso et al., 2015), esophagus (GSE97687)(Najgebauer et al., 2019), prostate (GSE115413 and GSE86258) (Lawrence et al., 2020), and stomach cancer (GSE117087, GSE194259 and GSE97686) (Maeda et al., 2020; Najgebauer et al., 2019). The samples clustered primarily according to the tissue in dimensions 1 and 2 (A), whereas they were separated into NAFs and CAFs by the fourth dimension (B). (C) Heatmap of 36 hypomethylated and 4 hypermethylated sites in CAFs versus NAFs, which were significantly differentially methylated in at least 4 of the tissues/cancers (mean methylation difference between NAFs and CAFs >10%; limma adjusted p-values <0.05). (D) Box plots of DNA methylation levels for all samples for the four differently methylated hypomethylated sites shared by all five tissues (adjusted p values are based on the limma differential methylation analysis).

Subsequently, we performed differential methylation analysis between NAFs and CAFs for each cancer type separately. CpGs were selected where the difference in mean DNAm > 10% and the limma adjusted p-values was < 0.05. Significant hypo- and hypermethylation was found for 1,134 and 559 CpGs in liver, 923 and 596 CpGs in lung, 9,631 and 5,199 CpGs in prostate, 1,389 and 210 CpGs in esophagus, and 3,286 and 2,988 CpGs in stomach, respectively. Of note, 34 hypomethylated and four hypermethylated sites were shared by at least four of these cancer categories (Figure 4C, Supplemental Figure S5). Furthermore, two hypomethylated CpGs were found in common in all datasets: cg07134930 in histondeacetylase 4 (*HDAC4*), and cg05935904 (not related to a gene; Figure 4D).

As previously mentioned, *EDARADD* has been described as one of the most differentially regions in two independent studies of lung and prostate cancer CAFs (Lawrence et al., 2020; Vizoso et al., 2015). Notably, the same CpG site cg09809672 (*EDARADD*) was significantly differently methylated in all our comparisons except for esophagus. Moreover, this CpG site has also been described as one of the genomic sites with conspicuous age-associated DNA methylation changes (Bocklandt et al., 2011). Therefore, we investigated whether CAF-associated CpGs are generally age-associated, but this was not observed for cg07134930 and cg05935904. When we used three different epigenetic clocks on CAFs and NAFs from liver (Horvath, 2013; Horvath et al., 2018; Levine et al., 2018), there were no significant differences in epigenetic-age predictions, which was in line with previous reports for prostate CAFs (Lawrence et al., 2020). Overall, our integrative analysis showed that CAFs from different tissues have overlapping epigenetic features.

### DNA methylation at CAF-associated CpGs is indicative for prognosis

The prevalence of cancer-associated fibroblasts might be associated with adverse prognosis in various types of cancer (Ping et al., 2021). We have therefore investigated if the DNA methylation levels in the CAF-associated CpGs cg09809672 (*EDARADD*), cg07134930 (*HDAC4*), and cg05935904 were indicative for overall survival. Our analysis encompassed 32 datasets from diverse cancer types within The Cancer Genome Atlas project (TCGA). To estimate association with overall survival we initially performed Cox proportional hazards models, incorporating age and gender as variables. In five cancer types - liver hepatocellular carcinoma (LIHC), kidney renal clear cell carcinoma (KIRC), kidney renal papillary cell carcinoma (KIRP), low grade glioma (LGG), and uveal melanoma (UVM) – at least one of the three CpGs revealed a significant association with survival (Supplemental Table 3). Furthermore, the association between the DNAm profiles of the three CpGs with survival was underscored by Kaplan-Meier curves and log-rank test, comparing the 25^th^ percentile of patients with the lowest methylation at these sites with the remaining patients (Figure 5).

**Figure 5:**
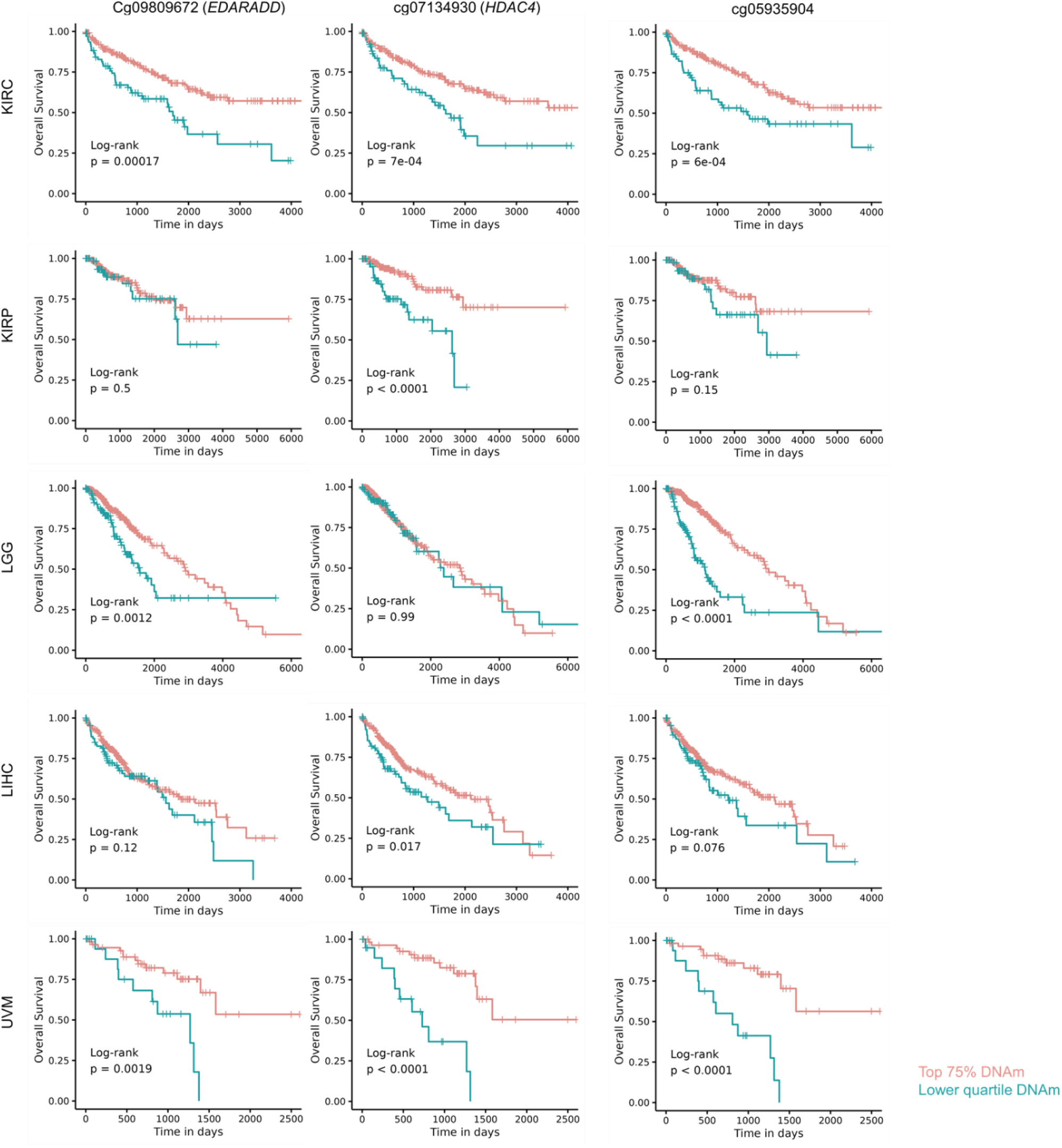
DNA methylation of cancer-associated fibroblasts is indicative for overall survival. Kaplan-Meier plots with overall survival for TCGA DNA methylation data of five different cancers: kidney renal clear cell carcinoma (KIRC), kidney renal papillary cell carcinoma (KIRP), low grade glioma (LGG), liver hepatocellular carcinoma (LIHC), and uveal melanoma (UVM). Patients were stratified by the 25^th^ percentile of lowest DNA methylation at the three CAF-associated CpGs (cg09809672 in *EDARADD*, cg07134930 in *HDAC4*, and cg05935904 without gene-association). Hypomethylation at these CpGs seems to be associated with higher CAF-content and shorter overall-survival.

## Discussion

Our results provide further evidence that interaction of cancer cells with their environment systematically remodels the epigenetic makeup of CAFs. Despite the heterogeneity of CAFs within a given tumor, among diverse tissues, and across individuals, there were consistent DNA methylation changes observed in multiple independent datasets. While not all liver-CAF samples exhibited distinct signatures, it is conceivable that the CAF^low^ samples encompass either less activated fibroblasts or distinct subpopulations. Previous research has identified different CAF-clusters (Yu et al., 2022; Zou et al., 2022) and it was even speculated that CAF subpopulations expressing different biomarkers might either promote or counteract tumor growth (Han et al., 2020; Zhao et al., 2023). In addition, Ma et al. have recently delved into the heterogeneity of CAF subsets using single cell gene expression data across different cancer types (Ma et al., 2023). It is conceivable that such diversity in CAF-subsets may also manifest in their methylome at the single-cell level, warranting further investigation in future studies. Either way, the CAF^low^ fraction revealed similar DNAm and transcriptomic patterns as NAFs and may thus not resemble a distinct cell type. Furthermore, analysis of methylome and transcriptome data consistently identified CAF^high^ samples with abnormal expression, particularly in candidate CAF-markers such as *RUNX3* (Koyama et al., 2023) and *EDARADD* (Lawrence et al., 2020). The heterogeneity in CAFs in transcriptome and DNAm might also originate from different cellular sources recruited into the CAF compartment (Zhao et al., 2023). Nevertheless, our analysis suggests that CAF-associated DNAm profiles closely resemble those of normal fibroblasts, hinting at a fibroblastoid cellular origin.

The discovery of reliable biomarkers for CAFs holds the potential to assist patient stratification and may ultimately provide new targets for therapeutic approaches (Han et al., 2020). In this study, we employed our CimpleG pipeline as a proof of concept to identify candidate CpGs for CAFs in liver tumors. Notably, hypomethylation at these CpGs was particularly observed in CAFs, to a lesser degree in NAFs, and exhibited high methylation levels in all other cell types examined. While our study establishes an association between hypomethylation at three of the eight tested CpGs and shorter overall survival in liver cancer, it is important to note that these biomarkers were selected based on a relatively small set of CAF samples. Consequently, further validation across independent cohorts is warranted in future studies.

It was remarkable to observe that apparently similar genomic regions become hypomethylated in CAFs across various studies and cancer types. Specifically, the three tested CpGs (cg09809672 in *EDARADD*, cg07134930 in *HDAC4*, and cg05935904) exhibited very consistent hypomethylation in CAFs compared to NAFs across all studies examined. In a prior study, Zou and coworkers focused on gene expression data to identify a signature of seven genes that were highly expressed in fibroblasts, and upregulated in ovarian cancer stroma compared with normal ovarian stroma (Zou et al., 2022). Notably, their candidate genes comprised *COL1A1*, which featured one of the most prominent hypomethylated regions in our CAF^high^ fractions, and podoplanin (*PDNP*), showing differential methylation in CAF^high^ *versus* NAFs. Their research linked elevated expression of this CAF signature with unfavorable prognosis in various cancer types. Our findings align with this association, revealing a similar link between hypomethylation at the three identified CpGs in CAFs and adverse outcomes across up to five different cancer types, with particularly pronounced effect in kidney renal clear cell carcinoma (KIRC).

In summary, our analysis highlights aberrant DNAm patterns in liver associated CAFs, demonstrating a clear association with differential gene expression. Moreover, these DNAm patterns exhibit notable overlap with CAF-signatures identified in other studies across various cancer types. This consistency suggests that there are reliable and reproducible epigenetic modifications occurring during the activation of CAFs, providing a discernable trace at the DNA methylation level.

## Experimental Procedures

### Cell isolation and cell culture

Tissue biopsies from liver tumors and adjacent healthy tissues were received from the Clinic for General, Visceral, and Transplantation Surgery at RWTH Aachen after written consent and following the guidelines of the Ethic Committee for the Use of Human Subjects at the University of Aachen (Permit number: EK 206/09). Tissue pieces were minced into small pieces, washed with PBS and incubated at 37°C in collagenase IV (1 mg/ml) containing medium (KnockOut DMEM from Gibco) overnight. After filtering through a strainer, the cells were cultured in DMEM medium containing 10% human platelet lysate, Penicillin-Streptomycin (100 U/ml), and L-Glutamine (2 mM). Cells were expanded at 37°C and 5% CO_2_ for 2-3 passages upon reaching confluence. CAFs and NAFs were successfully isolated from 11 patients (5 hepatocellular carcinoma [HCC], 5 liver metastasis of colon cancer [CRLM] and 1 liver metastasis of anal cancer [ALM]; mean age 62.6 ± 12.8 sd; 8 male and 3 female).

### Immunostaining

For the immunostaining, cells were cultured on gelatin (0.1%) coated cover slips and fixed with 2-4% PFA for 15 min. Cells were permeabilized for 30-60 min with 0.5% TWEEN 20 or 0.1% Triton™ X in PBS containing 5% BSA. Primary antibodies for vimentin (Sigma-Aldrich), α-SMA (Sigma-Aldrich) and pan-cytokeratin (Sigma-Aldrich) were added overnight at 4°C (Supplemental Table S4). The next day secondary antibodies Alexa 594 and Alexa 647 (ThermoFisher) were added for 1 hour. Nuclei were counter-stained with DAPI.

### Flow cytometry

Cells were fixed in 2% PFA for 15 min and stained with conjugated antibodies for 30 min. These included mouse-anti-human antibodies for CD14, CD29, CD31, CD34, CD45, CD73, CD90, and CD105 (Supplemental Table S4). Afterwards, cells were kept in PBS with 2% FCS. Samples were measured with a FACSCanto II (BD Biosciences) and the FlowJo software was used to analyze the data.

### DNA methylation analysis

Genomic DNA was isolated with the NucleoSpin Tissue kit (MACHEREY-NAGEL) and hybridized to Illumina MethylationEPIC BeadChips (at Life and Brain, Bonn, Germany). Initial quality control of DNA methylation data was performed with the minfi package (v1.48.0), and three samples with low overall signal intensities were removed at this step. The SeSAMe package (v1.20.0) (Zhou et al., 2018) was used for preprocessing (“QCDPBG”) including dye bias correction, quality mask filtering (Zhou et al., 2016), NOOB normalization (Triche et al., 2013) and calculation of detection p-values. CpG probes, which failed in 10% or more samples, non-cg probes, probes on X- and Y-chromosomes or probes flagged in the b5 manifest were removed. The data was then converted into a GenomicRatioSet to apply minfi-based functions. Additionally, we removed two outlier NAF samples.

For the public data from GEO (Supplemental Table S2) .idat files were preprocessed in the same way. The data was preprocessed separately as three different datasets and then merged. If no .idat files were available we utilized signal intensities and beta matrices. The public data consisted of Illumina 450K and EPIC data and was therefore reduced to the probes that overlap.

Limma (v.3.58.0) was used for the multidimensional scaling (MDS) plots and the differential methylation analysis, where probes with ≥ 0.2 difference in mean beta values and adjusted p-values (Benjamini-Hochberg) ≤ 0.05 were considered significant. The ComplexHeatmap package (v2.18.0) was used to generate the heatmaps including Pearson correlation as distance and the “ward.D2” method for clustering. Gene ontology analysis was done with the missMethyl R package (v.1.36.0). The DMRcate package (v2.14.1) was used to identify differently methylated regions, which were defined as regions of 1000 bp that included at least 2 differentially methylated CpGs with a “betacutoff” of 0.2 and a “pcutoff” of FDR = 0.05. The UpSetR package (v1.4.0) was used to make the upset plots. The DMR methylation plots were done using the Gviz (v1.46.0) and org.Hs.eg.db R packages (v3.18.0). To calculate the epigenetic age of the samples the wateRmelon R package (v2.8.0) was used.

For selection of top ranked CAF-specific CpGs we used our previously developed R package CimpleG (v0.0.5.9001) (Maie et al., 2023). For this purpose, we divided the samples randomly into a selection and test set based on an 80/20 split. We also performed a pre-selection of CpGs based on at least 20% mean difference in methylation values between all NAFs and CAFs.

### RNA-seq analysis

Total RNA was isolated using the NucleoSpin RNA kit (Macherey-Nagel). For library preparation the TruSeq Stranded mRNA kit (Illumina) was selected and sequenced on a NextSeq 500 (Illumina) using the NextSeq 500/550 High Output Kit v2.5 (150 cycles). Sequencing was performed at IZKF-associated Genomics Facility of the RWTH University. The nf-core/rnaseq pipeline was applied for alignment using STAR (hg38 genome) and generation of the count matrix using Salmon. Analysis was done with DESeq2 in R (Love et al., 2014). Generally, genes with overall less than 10 counts were removed. Genes were considered significantly differentially expressed when they showed a log2 fold change ≥ 2 and adjusted p-value ≤ 0.1. For visualization the data was VST transformed. RNA-seq and DNA methylation data was matched and combined by annotating both to Ensemble gene IDs.

### Survival analysis

Data gathering, curation and analysis was performed in R. Data for all TCGA projects, was downloaded with the TCGAbiolinks R package (v2.28.3), setting the parameters data.category to “DNA Methylation”, sample.type to “Primary Tumor”, platform to “Illumina Human Methylation 450” and data.type to “Methylation Beta Value”. Any sample that did not have information for the CpGs under analysis was dropped. Focusing on the clinical data, samples that showed up as duplicated or for which the survival clinical variables (deceased, days_to_death and days_to_last_follow_up) could not be found were not considered. For the Kaplan-Meier plots and log-rank tests, cancer patients were stratified by the 25^th^ percentile of DNAm at the respective CpG. For the Cox proportional hazards models, the continuous methylation value was used for each CpG (cg09809672, cg07134930, cg05935904) with gender and age as additional model variables. The p-value associated to each model variable was corrected using Benjamini and Hochberg p-value correction procedure.

## Further Information

### Data availability

All raw data are available from the corresponding authors upon reasonable request. The accession IDs of all public datasets used are mentioned and referenced in Supplemental Table S2.

### Author Contributions

M.S. performed the experiments as well as the analysis of methylation and gene expression data. T.M. performed survival analysis. W.W. and I.C. initiated the research, and W.W. designed and supervised the study. M.S., T.M., and W.W. wrote the manuscript, and all authors approved the final version.

### Conflicts of Interest

W.W. is involved in the company Cygenia GmbH (www.cygenia.com) that can provide service for epigenetic analysis to other scientists. Apart from this, the authors have no competing interests to declare.

### Funding

This research was supported by the Deutsche Forschungsgemeinschaft (DFG: 363055819/GRK2415 (W.W.); WA1706/11-1 (W.W.); WA 1706/12-2 (W.W.) and GE 2811/4-1 (I.C.) within CRU344/417911533; WA1706/14-1 (W.W.); and SFB 1506/1 (W.W.)); the ForTra gGmbH für Forschungstransfer der Else Kröner-Fresenius-Stiftung (W.W.), the BMBF Fibromap Consortia (I.C.), and particularly by the Interdisciplinary Center for Clinical Research within the faculty of Medicine at the RWTH Aachen University (W.W. and I.C.: IZKF O3–3).

## Acknowledgements

We want to thank Mohamed Mabrouk who supported alignment and quantification of RNA-sequencing data.

