## Supplemental Figures S1-S5, Table S3 & S4 for "Cancer-associated fibroblasts reveal aberrant DNA methylation across different types of cancer"

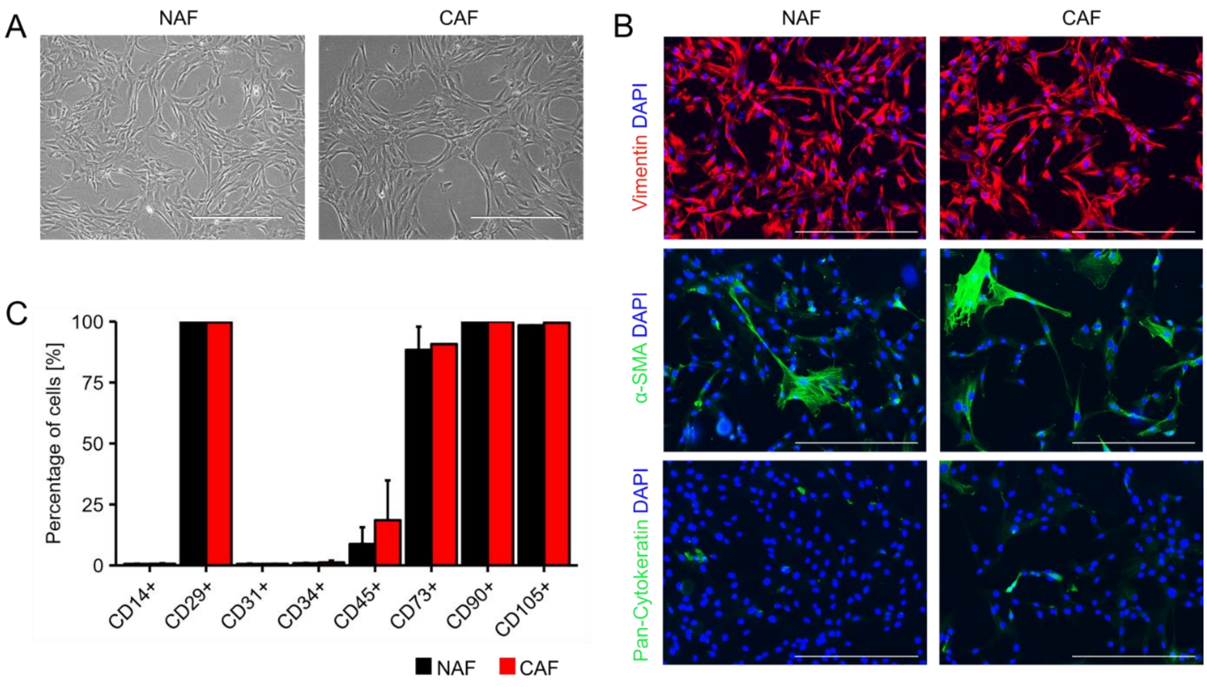


**Figure S1: Characterization of CAFs from liver (related to Figure 1)**

(A) Phase contrast images of normal tissue-associated fibroblasts (NAFs) and cancer-associated fibroblasts (CAFs) from one exemplary donor. Scale bar = 400 µm.

(B) Immunostaining of NAFs and CAFs from one exemplary donor. Stained with antibodies for vimentin, alpha smooth muscle actin (αSMA), and pan-cytokeratin. Scale bar = 400 µm.

(C) Flow cytometry results across different donors (n = 9). All cell preparations represent a typical fibroblastoid immunophenotype and are positive for CD105, CD29, CD73 and CD90.

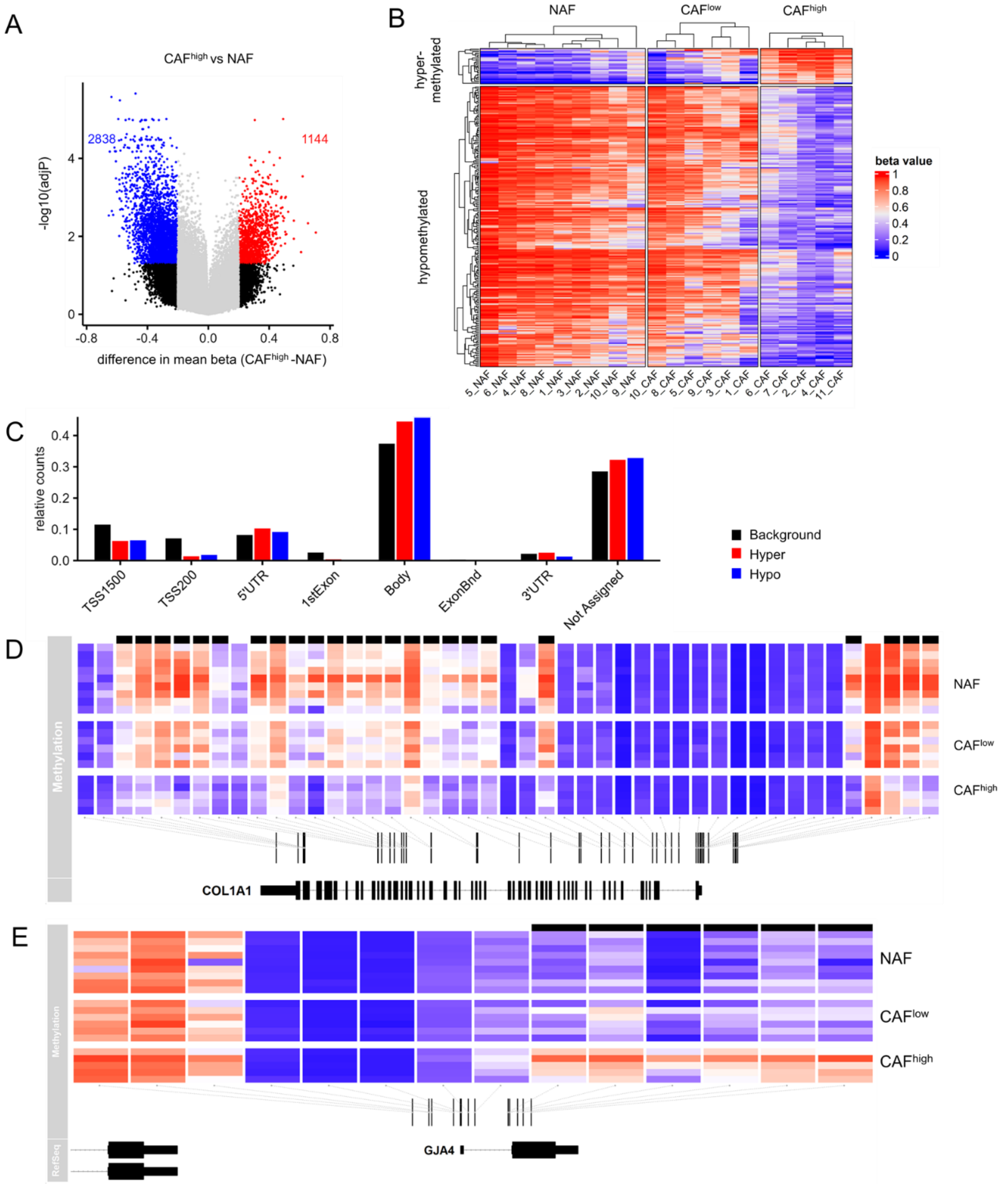


**Figure S2: Differential DNAm between CAF$^{high}$ and NAFs from liver tumors (related to Figure 1)**

(A) Volcano plot comparing methylation data (beta values) from NAF and CAF$^{high}$. Adjusted p-values from limma are depicted.

(B) Heatmap of the top 250 differently methylated CpG sites with the highest difference in mean beta values between CAF$^{high}$ and NAFs (limma adjusted p-values < 0.05).

(C) Distribution of differently methylated CpG sites to gene regions based on Illumina's annotation.

(D) Heatmap of DNAm of CpG sites associated with *COL1A1*. Black bars on top of the heatmap indicate significantly differently methylated sites between CAF$^{high}$ and NAFs. *COL1A1* contained the top differently hypomethylated region (DMR).

(E) The top differently hypermethylated region was found overlapping the gene *GJA4*.

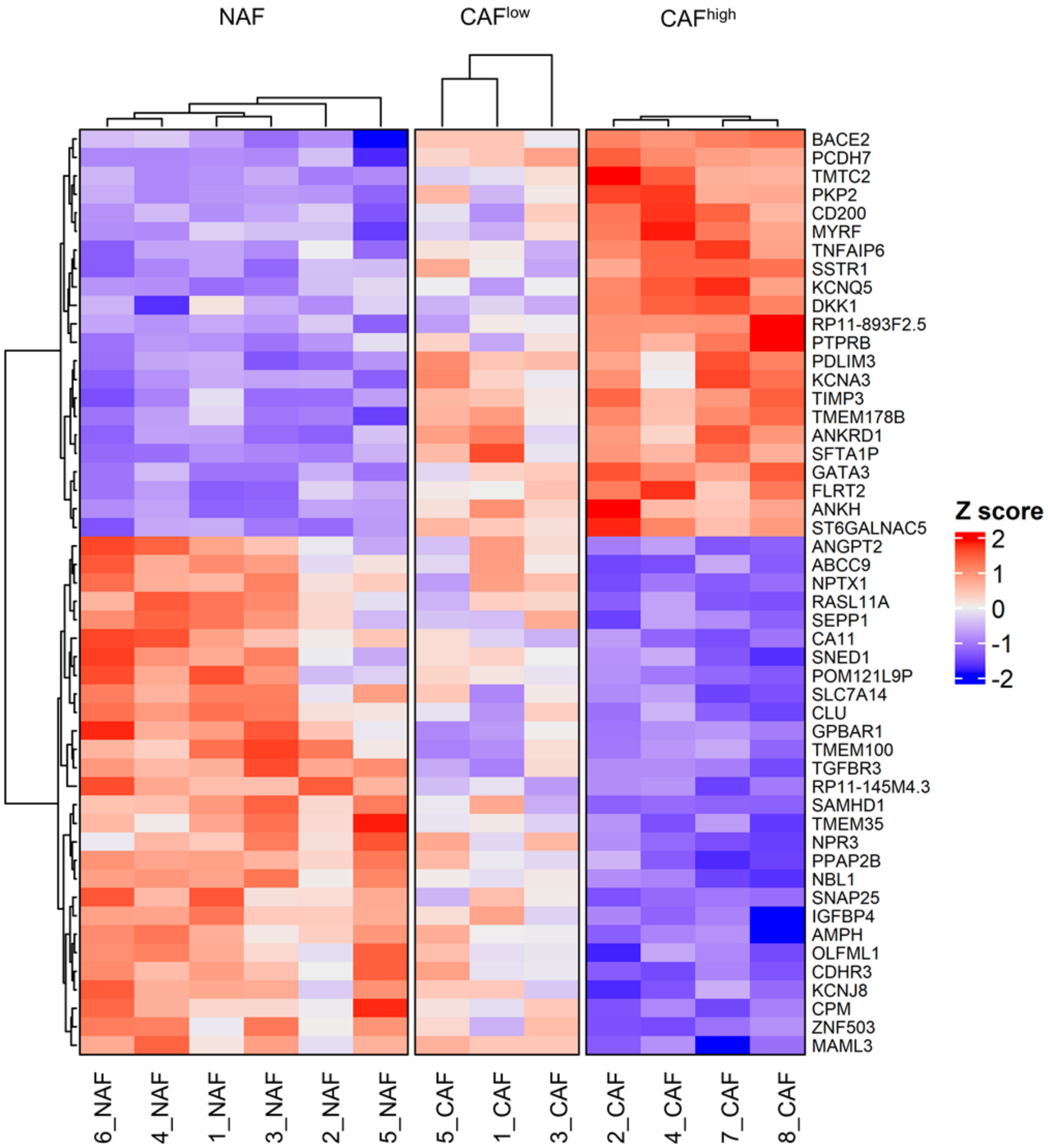


**Figure S3: Differentially expressed genes in liver CAFs (related to Figure 2)**

Top 50 most significantly different expressed genes between NAF and CAF$^{high}$. Depicted are Z scores of vst-transformed data from DEseq.

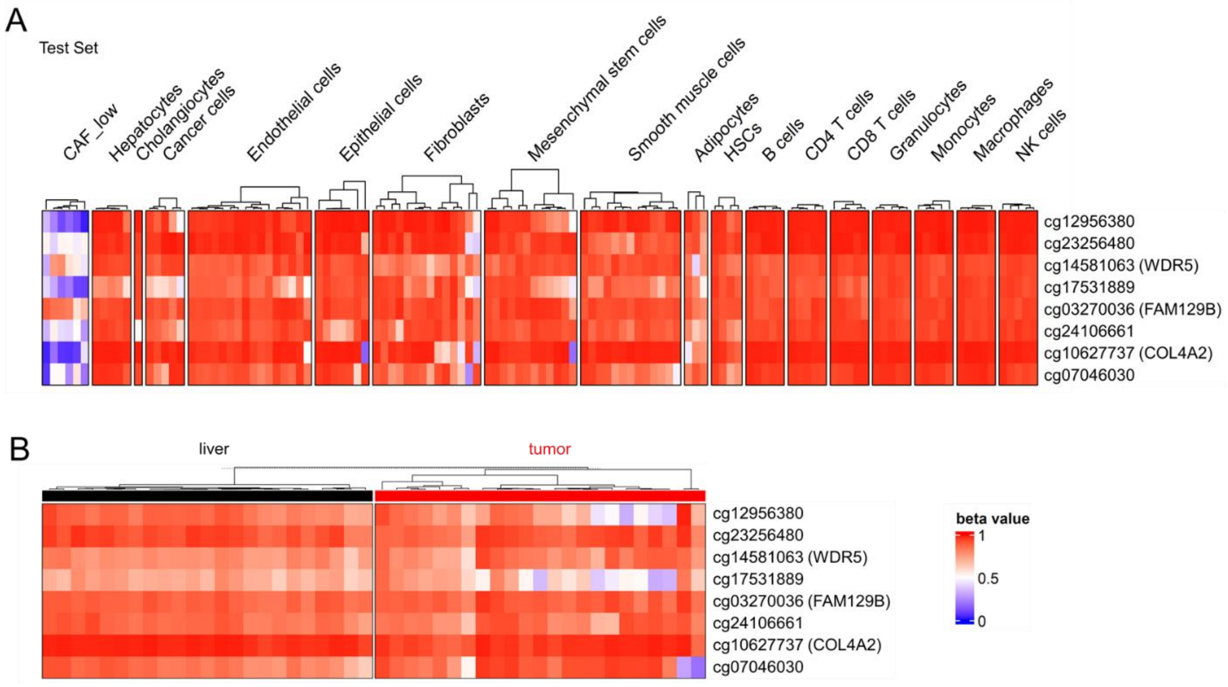


**Figure S4: Further benchmarking of DNA methylation biomarkers for CAFs in liver cancer. (related to Figure 3)**

(A) Heatmap of DNAm of eight candidate CpGs that were selected to discern CAFs from other cell types. The results of the test dataset are depicted here.

(B) Heatmap of DNAm of the eight candidate CpGs for CAFs in normal liver tissue as compared to hepatocellular cancer (GSE136380).

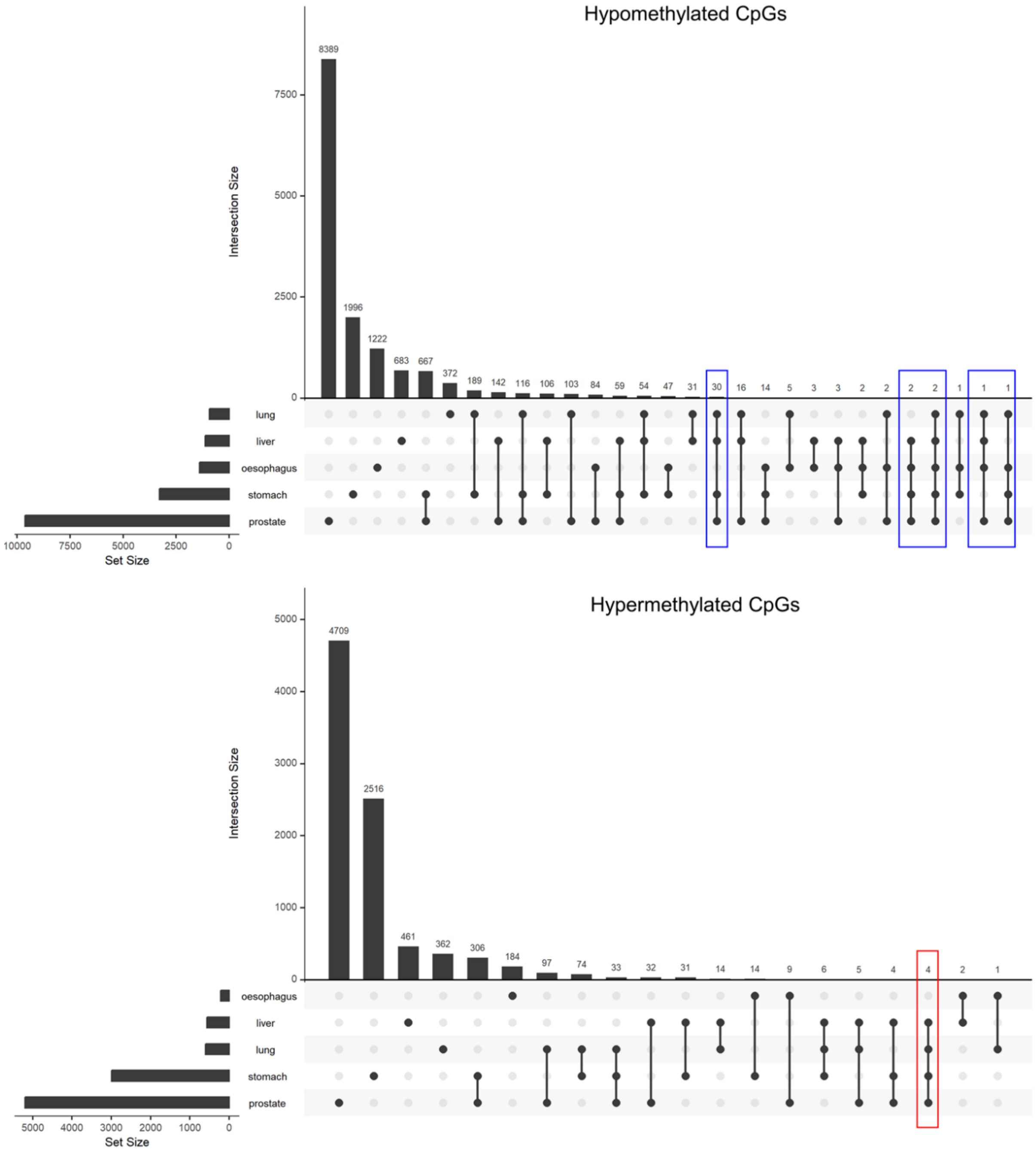


**Figure S5: Overlapping CAF-associated CpGs in different datasets (related to Figure 4)**

Upset plots of the overlap of differently methylated CpG sites between each of the different tissues and cancers. Differential methylation analysis was performed for each dataset independently. Columns that show the overlap of differently methylated sites for at least four groups are marked. This analysis was performed for hypo- and hypermethylated CpGs.

### Supplemental Tables

**Supplemental Table S3. Hazard ratios of COX regression models.**

| Cancer | CpG model | term | ci.95_lower | HR | ci.95_uper | adj_pval |
|---|---|---|---|---|---|---|
| KIRC | cg09809672 | gendermale | 0.72 | 1.08 | 1.62 | 0.92646 |
| | | age | 1.016 | 1.034 | 1.052 | 0.00258 |
| | | cg09809672 | 0.003 | 0.012 | 0.049 | < 0.00001 |
| | cg07134930 | gendermale | 0.741 | 1.11 | 1.665 | 0.85663 |
| | | age | 1.022 | 1.041 | 1.06 | 0.00021 |
| | | cg07134930 | 0 | 0.004 | 0.16 | 0.0294 |
| | cg05935904 | gendermale | 0.691 | 1.039 | 1.56 | 0.94981 |
| | | age | 1.016 | 1.034 | 1.053 | 0.00237 |
| | | cg05935904 | 0.015 | 0.066 | 0.296 | 0.00414 |
| KIRP | cg09809672 | gendermale | 0.269 | 0.528 | 1.037 | 0.24555 |
| | | age | 0.977 | 1.004 | 1.033 | 0.93486 |
| | | cg09809672 | 0.006 | 0.079 | 1.033 | 0.21173 |
| | cg07134930 | gendermale | 0.269 | 0.529 | 1.039 | 0.24555 |
| | | age | 0.981 | 1.008 | 1.035 | 0.82838 |
| | | cg07134930 | 0 | 0 | 0.007 | 0.00045 |
| | cg05935904 | gendermale | 0.256 | 0.503 | 0.987 | 0.19766 |
| | | age | 0.98 | 1.007 | 1.035 | 0.85663 |
| | | cg05935904 | 0.016 | 0.125 | 0.953 | 0.19766 |
| LGG | cg09809672 | gendermale | 0.714 | 1.021 | 1.461 | 0.95686 |
| | | age | 1.036 | 1.052 | 1.067 | < 0.00001 |
| | | cg09809672 | 0 | 0 | 0.013 | 0.00031 |
| | cg07134930 | gendermale | 0.746 | 1.067 | 1.525 | 0.92646 |
| | | age | 1.043 | 1.057 | 1.073 | < 0.00001 |
| | | cg07134930 | 0 | 0.039 | 6.719 | 0.46762 |
| | cg05935904 | gendermale | 0.775 | 1.111 | 1.591 | 0.82997 |
| | | age | 1.046 | 1.061 | 1.076 | < 0.00001 |
| | | cg05935904 | 0.002 | 0.007 | 0.031 | < 0.00001 |
| LIHC | cg09809672 | gendermale | 0.631 | 0.908 | 1.306 | 0.85089 |
| | | age | 0.996 | 1.01 | 1.024 | 0.42417 |
| | | cg09809672 | 0.048 | 0.182 | 0.692 | 0.07419 |
| | cg07134930 | gendermale | 0.603 | 0.868 | 1.25 | 0.75219 |
| | | age | 0.997 | 1.011 | 1.025 | 0.37921 |
| | | cg07134930 | 0.001 | 0.019 | 0.331 | 0.04297 |
| | cg05935904 | gendermale | 0.609 | 0.879 | 1.268 | 0.78258 |
| | | age | 0.998 | 1.012 | 1.026 | 0.33902 |
| | | cg05935904 | 0.401 | 0.983 | 2.413 | 0.98172 |
| UVM | cg09809672 | gendermale | 0.638 | 1.535 | 3.689 | 0.63071 |
| | | age | 1.009 | 1.05 | 1.093 | 0.08677 |
| | | cg09809672 | 0 | 0 | 0.003 | 0.00351 |
| | cg07134930 | gendermale | 0.592 | 1.408 | 3.348 | 0.74469 |
| | | age | 0.997 | 1.034 | 1.072 | 0.25712 |
| | | cg07134930 | 0.001 | 0.011 | 0.127 | 0.00339 |
| | cg05935904 | gendermale | 0.502 | 1.222 | 2.977 | 0.8984 |
| | | age | 1.018 | 1.061 | 1.107 | 0.03979 |
| | | cg05935904 | 0 | 0.002 | 0.044 | 0.00085 |

Hazard ratios (HR), 95% confidence intervals, and adjusted p-values are provided for all three relevant CpGs in five tumor models: KIRC = kidney renal clear cell carcinoma, KIRP = kidney renal papillary cell carcinoma, LGG = low grade glioma, LIHC = liver hepatocellular carcinoma, UVM = uveal melanoma. COX regression models were calculated for each cancer type and each of the three CpGs. Significant association with DNAm is highlighted in red.

**Supplemental Table S4: Antibodies used in the study**

| Antibodies | Label | Isotype | Clone | Manufacturer |
|---|---|---|---|---|
| Anti-human vimentin | - | Mouse IgM | LN-6, monoclonal | Sigma-Aldrich |
| Anti-human α-SMA | - | Mouse IgG2a | 1A4, monoclonal | Sigma-Aldrich |
| Anti-human pan-cytokeratin | - | Mouse IgG1/ IgG2a | C-11+PCK-26+CY-90+KS-1A3+M20+A53-B/A2, monoclonal | Sigma-Aldrich |
| Anti-mouse IgM Alexa 594 | - | Goat IgG | polyclonal | ThermoFisher |
| Anti-mouse IgG Alexa 647 | - | Goat IgG | polyclonal | ThermoFisher |
| Anti-human CD14 | APC | Mouse IgG2a | M5E2, monoclonal | BD Biosciences |
| Anti-human CD29 | PE | Mouse IgG1 | MAR4, monoclonal | BD Biosciences |
| Anti-human CD31 | PE | Mouse IgG1 | WM59, monoclonal | BD Biosciences |
| Anti-human CD34 | APC | Mouse IgG1 | 581, monoclonal | BD Biosciences |
| Anti-human CD45 | APC | Mouse IgG1 | HI30, monoclonal | BD Biosciences |
| Anti-human CD73 | PE | Mouse IgG1 | AD2, monoclonal | BD Biosciences |
| Anti-human CD90 | APC | Mouse IgG1 | 5E10, monoclonal | BD Biosciences |
| Anti-human CD105 | FITC | Mouse IgG2a | MEM-226, monoclonal | ImmunoTools |
